# HyMM: Hybrid method for disease-gene prediction by integrating multiscale module structures

**DOI:** 10.1101/2021.04.30.442111

**Authors:** Ju Xiang, Xiangmao Meng, Fang-Xiang Wu, Min Li

## Abstract

**Motivation:** Identifying disease-related genes is important for the study of human complex diseases. Module structures or community structures are ubiquitous in biological networks. Although the modular nature of human diseases can provide useful insights, the mining of information hidden in multiscale module structures has received less attention in disease-gene prediction.

**Results:** We propose a hybrid method, HyMM, to predict disease-related genes more effectively by integrating the information from multiscale module structures. HyMM consists of three key steps: extraction of multiscale modules, gene rankings based on multiscale modules and integration of multiple gene rankings. The statistical analysis of multiscale modules extracted by three multiscale-module-decomposition algorithms (MO, AS and HC) shows that the functional consistency of the modules gradually improves as the resolution increases. This suggests the existence of different levels of functional relationships in the multiscale modules, which may help reveal disease-gene associations. We display the effectiveness of multiscale module information in the disease-gene prediction and confirm the excellent performance of HyMM by 5-fold cross-validation and independent test. Specifically, HyMM with MO can more effectively enhance the ability of disease-gene prediction; HyMM (MO, RWR) and HyMM (MO, RWRH) are especially preferred due to their excellent comprehensive performance, and HyMM (AS, RWRH) is also good choice due to its local performance. We anticipate that this work could provide useful insights for disease-module analysis and disease-gene prediction based on multi-scale module structures.

**Availability:** https://github.com/xiangiu0208/HvMM

**Supplementary information:** Supplementary data are available at *Bioinformatics* online.

## 1 Introduction

The progress of human disease-gene discovery has helped understand the underlying molecular basis of human diseases, but genes known to be associated with diseases only account for a very small proportion of the incidences (Barabasi, et al., 2011; Kann, 2010; Moreau and Tranchevent, 2012; Wang, et al., 2011). Traditional approaches such as linkage analysis and genome-wide association studies often provide a long list of candidate genes, requiring expensive and time-consuming experimental identification (Botstein and Risch, 2003; Hirschhorn, 2009). Therefore, developing computational algorithms for predicting disease-related genes is indispensable to accelerate the discovery of disease-related genes (Li, et al., 2019; Luo, et al., 2019; Moreau and Tranchevent, 2012).

Human complex diseases can be recognized as the consequences of perturbations or functional abnormalities of biomolecule networks (del Sol, et al., 2010). Thus, network-based algorithms have been a popular strategy for disease-gene prediction (Cáceres and Paccanaro, 2019; Hu, et al., 2018; Lei and Zhang, 2019; Lin, et al., 2018; Luo and Liang, 2015; Moreau and Tranchevent, 2012; Peng, et al., 2017; Valdeolivas, et al., 2018; Yang, et al., 2015; Zeng, et al., 2017) and “guilt-by-association” provides a topdown central principle for predicting disease-related genes in the networks. For example, some algorithms infer disease-related genes by considering the shortest-path distance or closeness between candidate genes and known disease gene (set) in a network (Hsu, et al., 2011; Hu, et al., 2018; Zhu, et al., 2012); some algorithms make use of network propagation to extract disease-related information (Chen, et al., 2009; Cowen, et al., 2017; Jiang, 2015; Köhler, et al., 2008); some module-based algorithms are also applied to the analysis of disease genes/modules as well as related issues (Akram and Liao, 2017; Choobdar, et al., 2019; Kitsak, et al., 2016; Liu, et al., 2012; Sun, et al., 2011). As we know, module structures or community structures are ubiquitous in the biomolecule networks. The modular nature of human diseases can provide useful insights for the study of diseases, but it has not been fully explored for disease-gene prediction (Choobdar, et al., 2019; Oti and Brunner, 2007).

Generally, genes related to the same or similar diseases are more similar functionally and their products tend to be highly interconnected in biomolecule networks, forming so-called disease modules (Barabasi, et al., 2011; Dwivedi, et al., 2020). However, genes related to the same disease are usually found to be distributed in several modules/subnetworks by specific algorithms (Barabasi, et al., 2011; Lage, et al., 2007). There are several possible reasons for this phenomenon. (1) Complex diseases usually involve functional abnormalities of multiple genes, and these genes may have different functions, playing different roles in the development of complex diseases (Barabasi and Oltvai, 2004). (2) The existing biomolecule networks such as protein-protein interactions are still incomplete (Menche, et al., 2015; Peng, et al., 2016). This may cause the detected network modules to be broken and incomplete. (3) Detected modules in networks are often algorithm-specific, because specific definitions of modules are different for different module identification algorithms (Fortunato and Hric, 2016). For example, Disease Module Identification DREAM Challenge specifies a valid module size between 3 and 100 (Choobdar, et al., 2019). Furthermore, some module identification algorithms may split a large module into several small submodules, or aggregate several small modules into a large one, because of the resolution related to the intrinsic definition or mechanism of algorithms (Fortunato and Barthélemy, 2007; Xiang, et al., 2018; Xiang, et al., 2017). In this case, algorithms with flexible resolution may more effectively mine the module structures of networks.

Multi-scale structures widely exist in various natural and artificial complex networks, so multi-scale module detection is an important strategy for studying complex systems such as biomolecular networks (Ahn, et al., 2010; Mucha, et al., 2010). For example, a module in a protein network may contain several sub-modules, e.g., some protein complexes (such as SAGA) contain several secondary complexes. Most of biological information (e.g., gene function and disease phenotype data) is organized in the form of hierarchical structure. Dunn, et al. (2005) used edge-betweenness clustering to separate protein interaction networks into modules correlating to annotated gene functions, where modules of different sizes can be identified by removing different numbers of edges. Lewis, et al. (2010) investigated the correlation between the functions of sets of proteins and network module/community structures at multiple resolutions/scales, and they showed that there exist different important scales of module/community structures depending on studied proteins and processes. Wang, et al. (2011) proposed a fast hierarchical clustering algorithm using the local metric of edge clustering value, which can uncover the hierarchical organization of functional modules that approximately corresponds to the hierarchical structure of GO annotations. In the DREAM Challenge, extended (multiscale) modularity optimization was used to identify disease modules (Choobdar, et al., 2019). Multiscale module structures can provide more information than single-scale one, but there are still many challenging problems, such as how to mine the valuable information hidden in the multiscale structures.

In order to make use of multiscale module structures to predict disease-related genes more effectively, we propose a hybrid method integrating the information of multiscale modules (HyMM), which consists of three key stages: extraction of multiscale modules, disease-gene scorings/rankings based on multiscale module structures and integration of multiple gene rankings (see **Figure 1**). The rest of the paper is organized as follows. In section 2, we introduce three multiscale module decomposition algorithms, including the sampling of multiscale module partitions, and propose the method for gene scorings/rankings based on multiscale modules. Then, we present a theoretical framework for integrating multi-feature information from multiscale modules or others, based on naïve Bayes theory. In section 3, we first introduce the experimental settings. Then, we conduct statistical analysis of multiscale modules, display the effectiveness of disease-gene scorings based on multiscale modules in disease-gene prediction, and verify the performance of HyMM by 5-fold cross-validation and independent test. Finally, we come to a conclusion.

**Figure 1.**
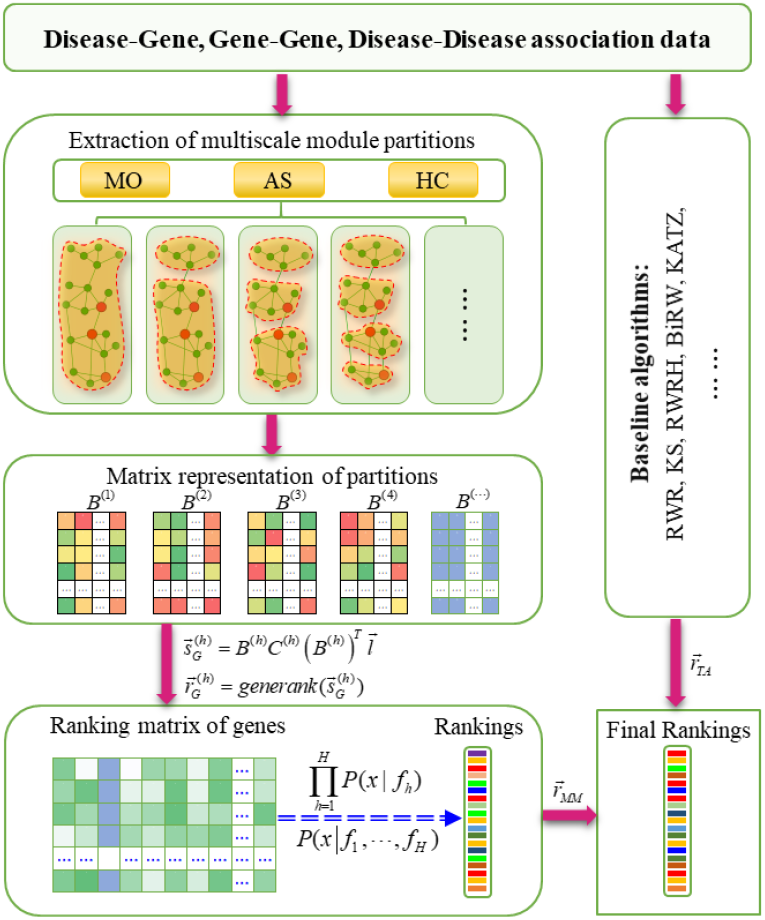
Workflow of the hybrid method for integrating the information of multiscale modules to predict disease-related genes in the network (HyMM).

## 2 Methods

Here, we propose a hybrid method for disease-gene prediction by integrating multiscale module structures (see **Figure 1** for the workflow of HyMM). In the following, we introduce the three key steps of HyMM: extracting multiscale modules, rankings of disease-related genes based on multiscale modules, and integration of multiple gene rankings.

### 2.1 Extraction of multiscale modules

Many algorithms have been proposed to detect module/community structures at different scales (Fortunato and Hric, 2016; Singhal, et al., 2020). We here use three typical multi-scale module identification algorithms (see **Figure 1**): modularity optimization (MO) (Newman and Girvan, 2004; Xiang, et al., 2015), asymptotic surprise (AS) (Xiang, et al., 2019) and fast hierarchical clustering (HC) (Wang, et al., 2011), all of which have flexible parameters to adjust the scales or sizes of modules. These algorithms can extract a set of network module partitions that contain important information of network structure, which can help predict disease genes.

#### 2.1.1 Multiscale module-identification algorithms

1. Modularity optimization (MO). MO detects module structure via optimizing modularity *Q*—a widely used quality function of module structure (Newman and Girvan, 2004). MO has been widely applied to module detection in networks (Choobdar, et al., 2019; Rahiminejad, et al., 2019). Given a module partition of a network, modularity *Q* can be calculated as,

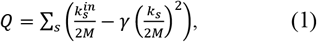

where *γ* is the resolution parameter; *M* is the total number of edges in the network; 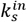 the inner degree of module *s; k_s_* the total degree of module s; the sum over all modules in the network. MO can detect module structures at different scales by adjusting the resolution parameter *γ*. It can find large-size modules if the *γ*-value is small. It can find small-size modules if the *γ*-value is large. Network can be decomposed into a set of single-node modules if *γ* is large enough, e.g., 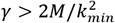 where *k_min_* is the minimum node degree in the network (Xiang, et al., 2012).
2. Asymptotic surprise (AS). It is an extension of surprise for community/module detection, originally proposed by Aldecoa, et al. (Aldecoa and Marín, 2014). Traag, et al. proposed the asymptotic approximation of surprise for weighted networks, by considering only the dominant term and using Stirling’s approximation of the binomial coefficient (Traag, et al., 2015). Recently, the multiscale version of the asymptotic surprise was proposed by adjusting the null model (Xiang, et al., 2019). It can be written as,

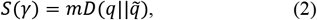

where *D*(*x||y*) = *x ln*(*x/y*) + (1 – *x*) *ln*((1 – *x*)/(1 – *y*)) is the Kull-back-Leibler divergence, which measures the distance between two probability distributions *x* and *y; q* = *m_int_/m* denotes the probability that an edge exists within a community; *m* is the number of existing edges in the network; while *m_int_* is the number of existing intra-community links in the partition; 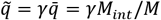 denotes the expected value of *q; M* is the maximal possible number of edges in a network; *M_int_* is the maximal possible number of intra-community edges in a given partition; γ is the resolution parameter.
3. Fast hierarchical clustering (HC). It is a fast hierarchical clustering algorithm based on the local metric of edge clustering value (ECV) (Wang, et al., 2011),

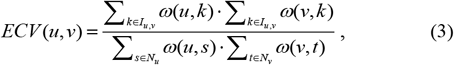

where *N_u_* and *N_v_* denote the sets of neighbors of nodes *u* and *v*, respectively; *I_u,v_* = *N_u_* ⋂ *N_v_* denotes the set of common neighbors of nodes *u* and *V; ω*(*u, s*) denotes the weight of edge between *u* and *s*. Given an undirected network and a threshold *λ,* a subgraph H is a *λ*-module if

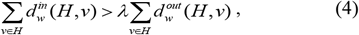

where *λ* is a tunable parameter; 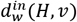 and 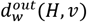 denote the in- and out-degrees of *H,* respectively. HC is an agglomerate algorithm. First, all nodes in the network are initialized as singleton modules. Then, all the edges are enqueued into a queue in nonincreasing order in terms of their ECV-values. The higher ECV-value the edge has, the more likely its two nodes are inside a module. Finally, HC assembles all the singleton modules into *λ* -modules by gradually adding edges in the queue to modules. One can get *λ* -modules of different sizes in the network by adjusting the value of *λ*. Generally, the larger the *λ*-value is, the larger the size of modules and thus the less the number of modules. In order to make the correlation between tunable parameter and module sizes consistent with other algorithms, we define a threshold parameter *γ* equivalent to *λ* by,

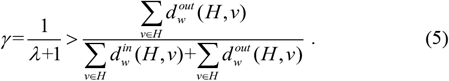 It provides a threshold for the ratio between out-degree and total degree of module.

#### 2.1.2 Sampling of multiscale module partitions

To extract a set of meaningful module partitions at different scales {*Ψ*_1_, *Ψ*_2_, … *Ψ_h_*, … *Ψ_H_*}, we first define a meaningful range [*γ_min_, γ_max_*] of *γ*-values, which covers all possible resolution samplings. Theoretically, *γ_min_* and *γ_max_* can be defined as *γ_min_* = *max*{*γ*|#[*Ψ_h_*] = 1]} and *γ_max_* = *min*{*γ*|#[*Ψ_h_*] = *N*]}. Here, we use the semi-empirical boundary for the three multiscale algorithms mentioned above, because it is usually difficult to obtain accurate interval boundaries.

Then, we use exponential sampling to sample *γ*-values from *γ_min_* to *γ_max_*, because it can give a reasonable coverage to different scales in the network, according to previous research (Xiang, et al., 2015; Xiang, et al., 2019). The exponential sampling uses a set of *γ* -values that are equally spaced on a logarithmic scale, i.e.,

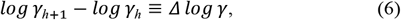

Here, *Δ log γ*=0.01 for default. The multiscale methods can extract a set of module partitions corresponding to the set of sampled *γ*-values.

### 2.2 Rankings of genes based on multiscale module structures

Given the set of extracted module partitions {*Ψ*_1_, *Ψ*_2_, … *Ψ_h_*, … *Ψ_H_*}, known disease-gene associations as well as disease-disease associations, we calculate the disease-relatedness scorings of modules and genes for each module partition one by one, and then generate a disease-association scoring/ranking matrix of genes (see **Figure 1**). The hypothesis of evaluating the disease relatedness of modules and genes is that the larger the proportion of disease-related genes in a module, the more likely the module is related to the disease, and thus the more likely other genes in the module are related to the disease.

#### Definition 1

A vector indicating association scores between *N* genes and a disease under study is defined as,

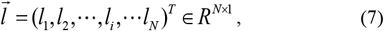

where *l_i_* = 1 if the *i*-th node is a known disease-related gene and *l_i_* =0 otherwise in PPI network.

#### Definition 2

A partition matrix *B*^(*h*)^ is defined to indicate the *h*-th module partition, where 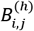 indicates whether gene *i* belongs to module *j* in this partition (see **Figure 1**).

#### Definition 3

A diagonal matrix *C*^(*h*)^ for each partition is defined as,

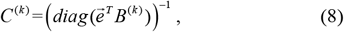

where 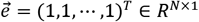.

Now, we evaluate the disease relatedness of all modules in the *h*-th module partition by,

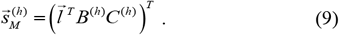

And the disease-relatedness scores of all genes can be calculated by,

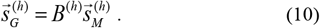

Then, we can introduce a scoring matrix of genes 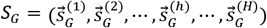 to record the gene scoring lists for all module partitions. Finally, we can transform the scoring matrix of genes into the ranking matrix of genes 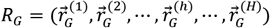 by the decreasing order of scores of genes,

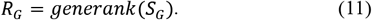

Note that the mean value of ranking is used for genes with the same scores.

According the above gene scoring/ranking strategy, genes within the same module have the same disease-relatedness scorings/rankings. Therefore, these scoring/ranking lists of genes contain the information of disease relatedness as well as the information of multiscale module structures from low to high resolutions. For example, if a module partition consists of two modules: one contains disease genes while not for another, gene scorings/rankings will have two values/levels. If the module with disease genes further split into two sub-modules with disease genes, then gene scorings/rankings will have at least three values/levels when there is no degeneracy of module scorings. The number of values/levels in the scoring/ranking list of genes is closely related to the number of disease-related modules in a module partition. As the resolution increases, we can get the gene scoring/ranking lists with more levels/values, thereby revealing different levels of disease-related information in the network.

#### Definition 4

From the perspective of kernel function, a kernel matrix for each module partition *Ψ_h_* is defined as,

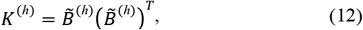

where 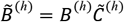 is a normalized partition matrix, 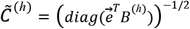 and 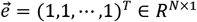.

This kernel matrix indicates the relationships between genes at the h-th partition, or say, the information extracted from this partition has been contained in the kernel matrix. As a result, all the information extracted from multi-scale module partitions can be recorded in the set of multiscale module kernel matrices {*K*^(*h*)^|*h* = 1~*H*}. And then, the disease-relatedness scores of all genes for module partition *Ψ_h_* can be calculated by,

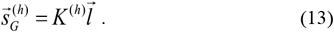

This provides another possible way to understand the scorings based on multi-scale module partitions.

### 2.3 Integration of multiple gene rankings

In order to effectively fuse the information derived from multiple module partitions, we introduce a theoretical framework for integrating multi-feature information based on naive bayesian theory. Given a set of features {*f_h_*|*h* = 1~*H*}, the probability at state *x* can be expressed as,

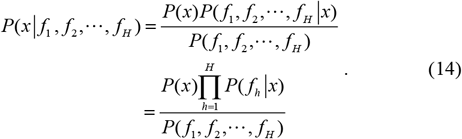

Here *P*(*f_h_|x*) = *P*(*f_h_*)*P*(*x|f_h_*)/*P*(*x*), where *P*(*f_h_*) denotes the prior probability of feature, and *P*(*x*) denotes the prior probability of a gene being at state *x*. Let *P*(*f_h_*) ≡ 1/*H* and *P*(*x*) ≡ *C*, then

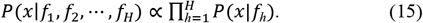

In disease-gene prediction, considering each module partition as a feature, we have calculated the scoring list 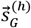 of candidate genes being related to a disease, and get the ranking list of the genes 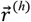 by the decreasing order of their scores. So, *P*(*x|f_h_*) can be defined as a function of the *h*-th scoring/ranking list of genes. Consider that the higher ranking of a gene means the larger value of the conditional probability *P*(*x_g_|f_h_*). Therefore, we can define 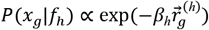 for each gene *g*. As a result, the above equation can be expressed as,

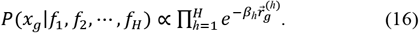

This study uses the strategy to get the comprehensive scores/rankings of all candidate genes for multiple features.

Given the above ranking matrix of genes from multiscale modules (MM), we calculate a comprehensive scoring list 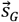 of genes and then get the ranking list of genes 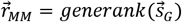 (see **Figure 1**).

The scorings/rankings from multiscale modules may provide useful and complementary information for disease-gene prediction, which is different from that of many other algorithms. So, we further integrate these rankings 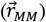 with that (denoted by 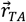) of eight typical algorithms (e.g., RWR, KS, RWRH and BiRW) (see **Figure 1**). The final scores/rankings of genes will be used to prioritize candidate genes. We will show that this integration can effectively improve the ability to predict disease genes due to information complementarity.

## 3 Results

In this section, we first introduce the experimental settings, including the datasets and evaluation methods. Then, we conduct statistical analysis of extracted multiscale modules, show the effectiveness of disease-gene scorings based on multiscale modules, and evaluate the performance of HyMM in predicting disease-related genes by a series of experiment tests and analyses.

### 3.1 Experimental settings

#### 3.1.1 Datasets

In order to conduct functional analysis of modules, we adopt three types of functional groups: Gene Ontology (GO) annotations, pathways, and disease-gene sets. To evaluate the predictive ability of algorithms, we employ the disease-gene associations, gene-gene associations and disease-disease associations. See **Supplementary Note 1** for details of datasets.

1. The GO annotations are downloaded from the Molecular Signatures Databases (MSigDB) (Consortium, 2018; Subramanian, et al., 2005). MSigDB omits GO terms with fewer than 5 genes or in very broad categories.
2. The pathway-gene sets are also obtained from the MSigDB, which are curated from several online pathway databases (such as KEGG and Reactome), publications in PubMed, and knowledge of domain experts (Kanehisa, et al., 2015; Matthews, et al., 2009).
3. Disease-gene sets. (a) We obtain the integrated disease-gene dataset (Ghiassian, et al., 2015; Menche, et al., 2015) retrieved from GWAS and OMIM (Hamosh, et al., 2005). MeSH is used to combine the different disease nomenclatures of the two sources into a single standard vocabulary. (b) Furthermore, we obtain the disease-gene associations from DisGeNet, and map the UMLS disease into MeSH disease (Piñero, et al., 2017). Dis-GeNET is known as a platform that contains one of the largest publicly available collections of disease-related genes.
4. The disease-disease associations are derived by using the associations between symptoms and diseases (Zhou, et al., 2014).
5. Gene-gene associations are derived from a comprehensive proteinprotein interaction (PPI) network that consists of the multiple sources of protein interactions (Menche, et al., 2015). The network data considers only physical protein interactions with experimental support. The identifiers of proteins are mapped into gene symbols.

Moreover, we construct disease-gene heterogenous network by integrating gene-gene associations, disease-gene associations and disease-disease associations mentioned above. Note that only the disease-gene associations in training set are used in the construction of the heterogenous network.

#### 3.1.2 Metrics for analysis of multiscale modules

In order to quantify the functional relevance of modules, we define the functional consistency of a module *M_m_* with respect to a functional group by, *G_f_* by,

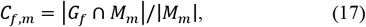

where a functional group denotes a set of genes with common characteristics and |*| denotes the number of elements in the group. For a set of functional groups, the functional consistency of a module is defined as the maximal functional consistency over all functional groups,

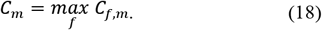

For a module partition *Ψ_h_*, consisting of a set of modules, its functional consistency is defined as the weighted average of the functional consistency scores of modules,

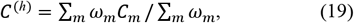

where *ω_m_* is proportional to the number of genes in the module.

#### 3.1.3 Evaluation methods

To investigate the performance of algorithms in disease-gene prediction, we use two evaluation strategies: traditional 5-fold cross-validation (5FCV) and independent test (IndTest).

1. For 5FCV, known disease-related genes for each disease are randomly split into five subsets. In each realization, one of the subsets is treated as a test set, while the rest is treated as a training set.
2. For IndTest, the disease-gene associations in the Mesh dataset are used as training set, and the disease-gene associations that belong to the DisGeNet dataset but do not belong to the Mesh dataset are used as test set.

Then, two kinds of standard evaluation metrics are used to quantify the performance of the prediction algorithms from the global and local perspectives respectively.

1. Global evaluation metric. The area under the receiver operating characteristic curve (AUROC) is a single scalar value and it is widely used to evaluate the comprehensive performance of the prediction algorithms. In a statistical sense, AUROC can be considered as the probability that the score of a randomly selected positive sample is larger than that of a randomly selected negative sample.
2. Local evaluation metric. Differing from AUROC, top-k Recall focuses on how many disease-related genes in the test set have been found in the top-k genes of the ranking list. As k-value varies, top-k Recall can provide a local performance curve intuitively to compare the performance of algorithms. For convenience of comparison, we further calculate the area under the top-k Recall curve (AURecall) to comprehensively and quantitatively evaluate the local performance of algorithms.

### 3.2 Statistical analysis of multiscale modules

We quantify the functional consistency of module partitions at different scales by using three functional groups: disease-gene set, pathways and GO annotations. The results shows that the functional consistency scores of module partitions at different levels increase with the increase of the resolution parameter (**Figure S1**). This means that the proportion of similar genes within the modules is increasing with the resolution parameter: these genes will more likely tend to have the same GO annotations, or belong to the same pathway or disease-gene set. This is because the edge density in the modules becomes higher with the increase of resolution parameter. Genes in the modules more likely tend to interact with each other, and thus have the same or similar functions or participate in common biological process. Therefore, the module partitions at different scales can provide different levels of information about the relationship between genes. This may provide more comprehensive understanding of genes and their functions.

Further, we analyze the distribution of disease-related genes/modules (referred as to modules containing disease-related genes) at different scales. As expected, the number of modules (*N*_m_) and disease-related modules (*N*_dm_) in module partition increases with the resolution parameter (see **Figure S2(a)-(c)**), because the network is being split into smaller and smaller modules. With the increase of resolution, the disease-related genes tend to be dispersed into more modules of smaller sizes (see **Figure S2(d)-(f)**), but these modules often have stronger functional consistency; at the same time, the total number of genes within the disease-related modules also becomes less and less (see **Figure S2(d)-(f)**), and the disease-related genes tend to be enriched in a subnetwork composed of the disease-related modules (see **Figure S2(g)-(i)**). As discussed above, the genes within these disease-related modules generally have stronger functional correlation (for diseases, pathways or GO). Thus, they are more likely to be disease-related genes.

### 3.3 Effectiveness of disease-gene scorings based on multiscale module structures

To check if multiscale module structures can really provide useful information for disease-gene prediction, we firstly use the comprehensive scorings of genes based on multiscale modules (i.e., HyMM(#, NULL)) directly to identify disease-related genes. The results show that HyMM(#, NULL), especially with MO, has comparable and even better performance in predicting disease-related genes (see **Figure 2, Figure 3, Figure 4, Figure 5**, and **Figures S3–S6**).

**Figure 2.**
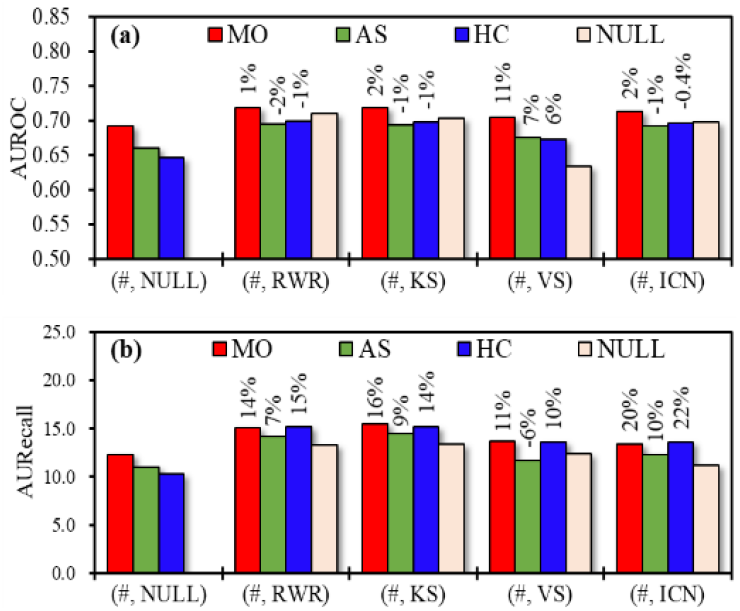
In the PPI network, performance comparison of (a) AUROC and (b) AURecall obtained by HyMM algorithms and four baseline algorithms. Multiscale module identification algorithms: MO, AS and HC. “(#, RWR)” denotes the HyMM algorithm integrating MO/AS/HC with RWR. “NULL” means that the multi-scale algorithm or baseline algorithm is not applicable. The percentage on the bar is performance improvement ratio to baseline algorithm.

**Figure 3.**
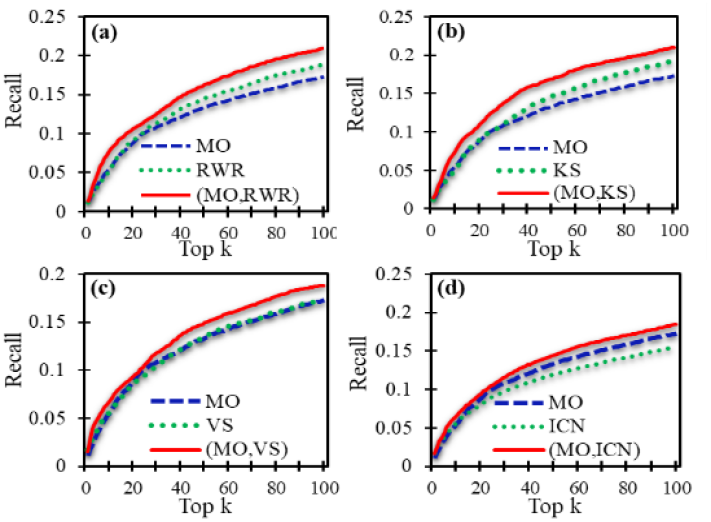
In the PPI network, top-k recall curves of HyMM with MO, compared to corresponding baseline algorithms: (a) RWR, (b) KS, (c) VS, (d) ICN.

**Figure 4.**
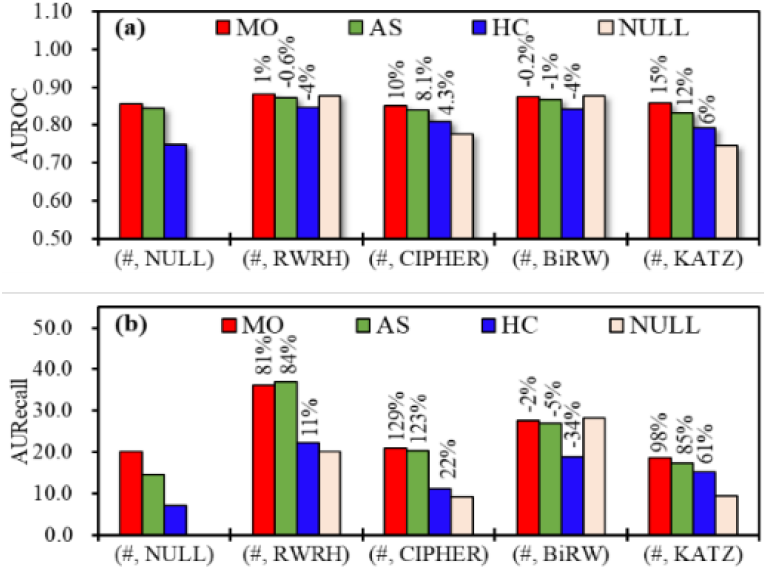
In the disease-gene heterogeneous network, performance comparison of (a) AUROC and (b) AURecall obtained by HyMM algorithms and four base-line algorithms: RWRH, CIPHER, BiRW and KATZ. “(#, RWRH)” denotes the HyMM algorithm with MO/AS/HC with RWRH. “NULL” means that the multi-scale algorithm or baseline algorithm is not applicable. The percentage on the bar is performance improvement ratio to baseline algorithm.

**Figure 5.**
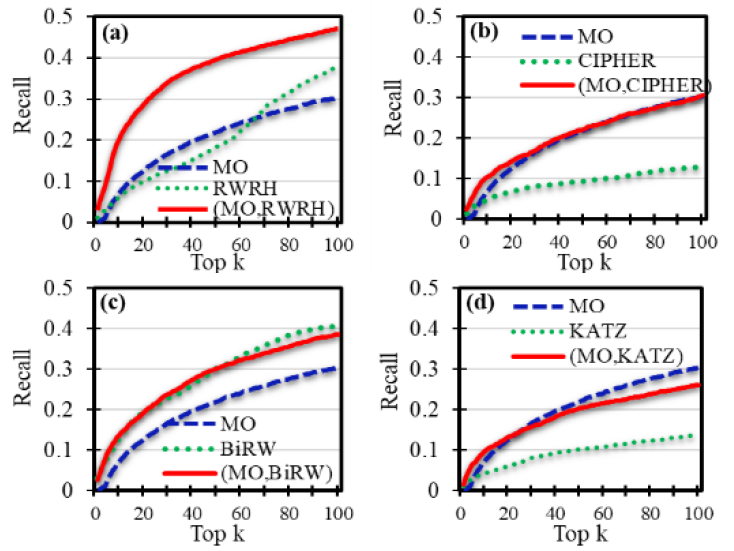
In the disease-gene heterogeneous network, top-k recall curves of HyMM with MO, compared to corresponding baseline algorithms: (a) RWRH, (b) CIPHER, (c) BiRW, (d) KATZ.

In the PPI network, for example, HyMM(MO/AS/HC, NULL) has larger AUROC-values than VS (**Figure 2**). The top-k recall curve of HyMM(MO, NULL) is similar to that of VS, and higher than that of ICN (**Figure 3**). The AURecall-values of the algorithms again confirm their above results of top-k recalls (**Figure 2, Figure 3** and **Figure S3–S4**). In the disease-gene heterogeneous network, HyMM(MO/AS, NULL) has larger AUROC- and AURecall-values than CIPHER and KATZ **(Figure 4** and **Figure 5**). Moreover, in terms of AUROC and AURecall (or top-k recall), the multiscale algorithm MO is better than AS and HC in diseasegene prediction.

### 3.4 Performance evaluation in cross-validation

The unique information from multiscale modules may improve the predictive ability of disease genes due to the information complementarity. Here, we first study the performance improvement of HyMM compared to eight baseline algorithms by 5-fold cross-validation, and then compare the performance of HyMM using different multiscale algorithms (see **Figure 2, Figure 3, Figure 4** and **Figure 5**)

#### 3.4.1 Performance improvement of HyMM to baseline algorithms

Here, we provide a pairwise comparison for each HyMM algorithm to the corresponding baseline algorithms both in the PPI network and the dis-ease-gene heterogeneous network.

In the PPI network, HyMM(MO,#), i.e., HyMM with MO, outperforms the corresponding baseline algorithms (RWR, KS, VS and ICN) both in terms of AUROC and AURecall (see **Figure 2**). In the disease-gene heterogeneous network, HyMM(MO,#) also outperforms the corresponding algorithms (RWRH, CIPHER and KATZ) both in terms of AUROC and AURecall, except for BiRW (see **Figure 4**); and HyMM with MO has the second highest AURecall improvement ratio for RWRH as well as the second highest AURecall-value, meaning that it has excellent local performance. These results indicate that the integration of MO can result in the performance improvement for most baseline algorithms, especially for the local performance.

HyMM with AS in the PPI network can improve the AURecall performance of RWR/KS/ICN, and only the AUROC performance of VS (see **Figure 2**). In the disease-gene heterogeneous network, HyMM with AS can improve the AURecall performance of RWRH/CIPHER/KATZ, and only the AUROC performance of CIPHER/KATZ (see **Figure 4**). And, HyMM with AS has the highest AURecall-improvement ratio for RWRH. These results means that the integration of AS also can lead to the large performance improvement in many cases, e.g., especially for the local performance of RWRH, though it is slightly inferior to MO in the global performance.

Also, HyMM with HC in the PPI network can only improve the AUROC performance of VS, but it can improve the AURecall performance of four baseline algorithms (RWR/KS/VS//ICN) (see **Figure 2**). HyMM with HC in the disease-gene heterogeneous network can improve the AURecall performance of RWRH/CIPHER/KATZ and the AUROC performance of CIPHER/KATZ, but the improvement ratio of HyMM with HC is inferior to HyMM with MO and AS (see **Figure 4**).

#### 3.4.2 Comparison and recommendation of HyMM using different multiscale algorithms

The above results show that, in most cases, HyMM with MO can improve the baseline algorithms more effectively than HyMM with other multiscale algorithms (AS and HC) (see **Figure 2** and **Figure 4**), obtaining better prediction performance, especially when integrating with RWRH in the heterogeneous network or integrating with RWR/KS in the PPI network.

The comparison between AS and HC shows that, in the PPI network, HyMM with HC has higher AURecall-values than HyMM with AS, while they have similar AUROC values (see **Figure 2**). In the disease-gene heterogeneous network, HyMM with AS has better AUROC and AURecall performance than HyMM with HC, and HyMM(AS, RWRH) has the highest AURecall-value (see **Figure 4**).

As a whole, based on the above analysis, the multiscale algorithm MO is preferred. In the PPI network, HyMM(MO,RWR) and HyMM(MO,KS) are both good choices because of their good global and local performance, while HyMM(HC, RWR/KS) is the second recommendation in terms of the local performance. In the heterogeneous network, HyMM(MO, RWRH) is the best recommendation in terms of its global and local performance, while HyMM(AS, RWRH) is the second recommendation in terms of its local performance.

### 3.5 Performance evaluation with external dataset

Furthermore, we calculate the scores of candidate genes by using the Mesh disease-gene associations as a training set, and then evaluate the prediction performance by using the disease-gene associations belonging only to Dis-GeNet as a test set.

In both the PPI network and the heterogeneous network, HyMM with MO can improve the global and local performance of all the baseline algorithms in pairwise comparisons (see **Figures S7–S8**). HyMM with AS/HC can improve the local performance of all the baseline algorithms in the PPI network as well as the baseline algorithms (except for BiRW) in the heterogeneous network.

In the PPI network, HyMM(MO,RWR) and HyMM(MO,KS) are still good choices in terms of global and local performance metrics, while HyMM(AS,RWR) and HyMM(AS,KS) are better candidates in terms of local performance metrics (see **Figures S7**). In the heterogeneous network, both HyMM(MO,RWRH) and HyMM(MO,BiRW) are good choices in terms of global and local performance metrics (see **Figures S8**).

As a whole, HyMM with MO has excellent performance in the independent test of external dataset; and HyMM(MO,RWR/KS) and HyMM(MO,RWRH) are still good recommendations.

### 3.6 Case study

As an example, we obtain a list of top-20 candidate genes for Alzheimer’s disease (AD) (see **Table S1**). Through literature verification, we find that many genes in the list have been proven to have associations with AD. For example, Park, et al. (2021) showed that ALK is important to the tau-mediated AD pathology; Pichiah, et al. (2020) showed that C4B was differentially expressed in AD; Stoye, et al. (2020) demonstrated that APOA1 might be a key factor within intestine altered in AD-like pathology. MTHFR and MAPT have been recorded as related to AD in DisGeNet. See **Supplementary Note 5** for details.

By the enrichment analysis of the above genes, we obtain the most relevant KEGG pathways and GO terms (see **Table S2** and **Table S3**), many of which have been known to be related to AD, such as the pathways: Lysosome, Metabolic pathways, Oxidative phosphorylation and Parkinson’s disease.

In addition, we analyze the druggability of the candidate genes (see **Table S1**), and find that many genes correspond to protein targets of approved or clinical trial-phase drug candidates (Finan, et al., 2017; Wang, et al., 2019) and many genes have a large number of interacted drugs (Freshour, et al., 2020), which may be potential therapeutic agents (see **Table S1**).

### 3.7 Parameter stability

HyMM mainly involves the sampling parameter, as well as the parameter 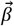 measuring multiple features. For simplicity, we set 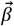 as a vector of equal weights, though optimizing the parameter can generate better results. We here study the effect of the sampling parameter *Δ log γ* on prediction performance, because it is closely related to the number of multi-scale module partitions and the amount of information extracted from the network.

We calculate the AUROC and AURecall values of all HyMM algorithms in the interval of *Δ log γ* ∈ [0.01,0.2] (**Figures S9–S10**). The results show that the HyMM algorithms with MO and AS are very stable to the sampling parameter, while the HyMM algorithms with HC have relatively large fluctuations.

## 4 Conclusion

Identifying human disease-related genes is important for the understanding of the underlying molecule basis of diseases and the prevention, diagnosis and treatment of human diseases. Network-based algorithms for disease-gene prediction are very popular, because human complex diseases are usually considered to be caused by the perturbations or functional abnormalities of disease-related modules or subnetworks in the biomolecule networks. However, multi-scale module structures are not fully utilized in the analysis and prediction of disease-related genes, though it widely exists in complex systems such as biomolecular networks. Therefore, we proposed the hybrid method called HyMM that integrates the information of multiscale modules to predict disease-related genes in the network, which can effectively enhance the ability of disease-gene prediction due to information complementarity.

Through three multiscale-module-decomposition algorithms (MO, AS and HC), we analyzed the functional consistency of multiscale modules and the distribution of disease-related genes in modules of different scales, and displayed the effectiveness of the information of multi-scale modules in identifying disease-related genes. By cross-validation and independent test, we confirmed the ability of HyMM to improve the performance of disease-gene prediction. The results showed that HyMM with MO can more effectively enhance the ability of disease-gene prediction in both global and local performances. HyMM(MO,RWR) and HyMM(MO,RWRH), as typical representatives, are especially preferred candidate algorithms, due to their excellent comprehensive performance in the PPI network and the heterogenous network, respectively. Moreover, HyMM(AS,RWRH) is also good choice due to its local performance.

Overall, this work provides an effective strategy for integrating multiscale module structures to enhance the ability of disease-gene prediction. We anticipate that it can provide useful insights for disease-module analysis and disease-gene prediction based on multi-scale module structures.

## Funding

This work was supported by the National Key Research and Development Program of China (Grant No. 2019YFA0706202), and the National Natural Science Foundation of China (Grant No. 61702054).

## Conflict of Interest

none declared.

## Supplementary Materials for

### Supplementary Note 1: Datasets

Here, we describe in detail the relevant datasets used in the study.

1. GO-gene sets. Gene Ontology (GO) is developed to provide biologically meaningful annotations of genes and their products (http://geneontology.org/). A GO annotation consists of a GO term associated with a specific reference that describes the work or analysis upon which the association between a specific GO term and gene product is based. The GO annotations here are downloaded from the Molecular Signatures Databases (MSigDB; http://www.gsea-msigdb.org/gsea/msigdb/collections.jsp), a collection of annotated gene sets (Consortium, 2018; Subramanian, et al., 2005). MSigDB omits GO terms with fewer than 5 genes as well as in very broad categories that would produce extremely large gene sets, and gene sets in each sub-collection are filtered to remove inter-set redundancy.
2. Pathway-gene sets. The pathway-gene sets here are also downloaded from the MSigDB (http://www.gsea-msigdb.org/gsea/msigdb/collections.jsp), which are curated from several online pathway databases (such as KEGG and Reactome), publications in PubMed, and knowledge of domain experts (Kanehisa, et al., 2015; Matthews, et al., 2009). These gene sets are often canonical representations of a biological process compiled by domain experts. As described in MSigDB, the pathway-gene sets are curated from the following online databases: BioCarta: http://cgap.nci.nih.gov/Pathways/BioCarta Pathways; KEGG: http://www.pathway.jp; Pathway Interaction Database: http://pid.nci.nih.gov; http://www.ndexbio.org; Reactome: http://www.reactome.org; SigmaAldrich: http://www.sigmaaldrich.com/life-science.html; Signaling Gateway: http://www.signaling-gateway.org; SuperArray SABiosciences: http://www.sabiosciences.com/ArrayList.php; WikiPathways: https://www.wikipathways.org/.
3. Disease-gene sets. We use two disease-gene datasets. (a) The first dataset is an integrated disease-gene dataset (Ghiassian, et al., 2015; Menche, et al., 2015) retrieved from GWAS (Genome-Wide Association Studies) and OMIM (Online Mendelian Inheritance in Man)(Hamosh, et al., 2005). We call it as the MeSH dataset, because MeSH (Medical Subject Headings ontology) is used to combine the different disease nomenclatures of the two sources into a single standard vocabulary. (b) The second dataset is obtained from DisGeNet (https://www.disgenet.org/), and we map the UMLS disease in the dataset into MeSH disease (Piñero, et al., 2017). DisGeNET is known as a platform that contains one of the largest publicly available collections of disease-related genes.
4. Disease-disease associations. The disease-disease association network is constructed by using the associations between symptoms and diseases. The strengths of these associations between symptoms and diseases are quantified through MeSH term co-occurrence, i.e., the number of PubMed identifiers where two Mesh terms occur together, and then they are normalized by the term frequency-inverse document frequency (Zhou, et al., 2014). Finally, the cosine similarity scores of the normalized symptom vectors of two diseases are used to quantify the association scores between the diseases.
5. Gene-gene associations. Because there exist noises in existing protein networks and the single-source protein networks are often incomplete, we here adopt a comprehensive protein interactome that consists of the following sources of protein interactions: regulatory interactions, binary interactions from several yeast two-hybrid high-throughput and literature-curated datasets, literature-curated interactions derived mostly from low-throughput experiments, metabolic enzyme-coupled interactions, protein complexes, kinase-substrate pairs and signaling interactions (Menche, et al., 2015). The network data considers only physical protein interactions with experimental support. The identifiers of proteins are mapped into gene symbols.

### Supplementary Note 2: Statistical analysis of multiscale modules

Here, we provide the results of statistical analysis of multiscale modules detected by using MO, AS and HC.

**Figure S 1.**
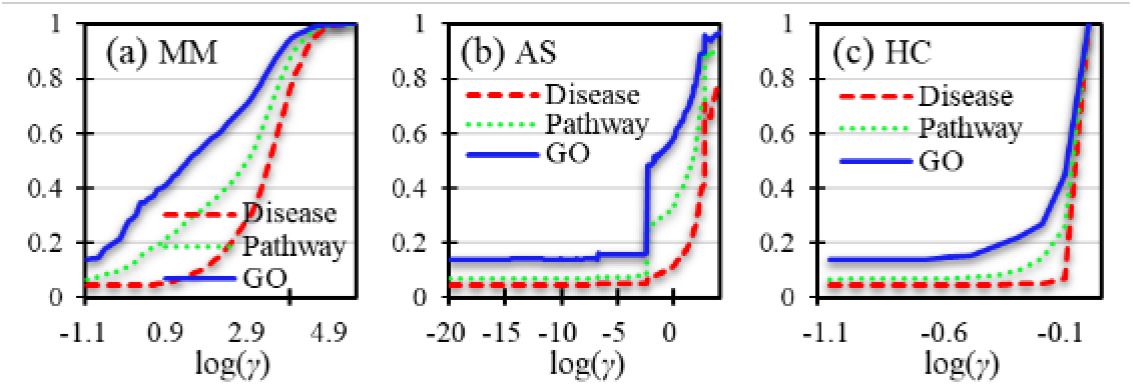
For three types of functional groups: GO annotations, pathways and diseases, functional consistency of module partitions detected by (a) MM, (b) AS and (c) HC, as a function of resolution parameter.

**Figure S 2.**
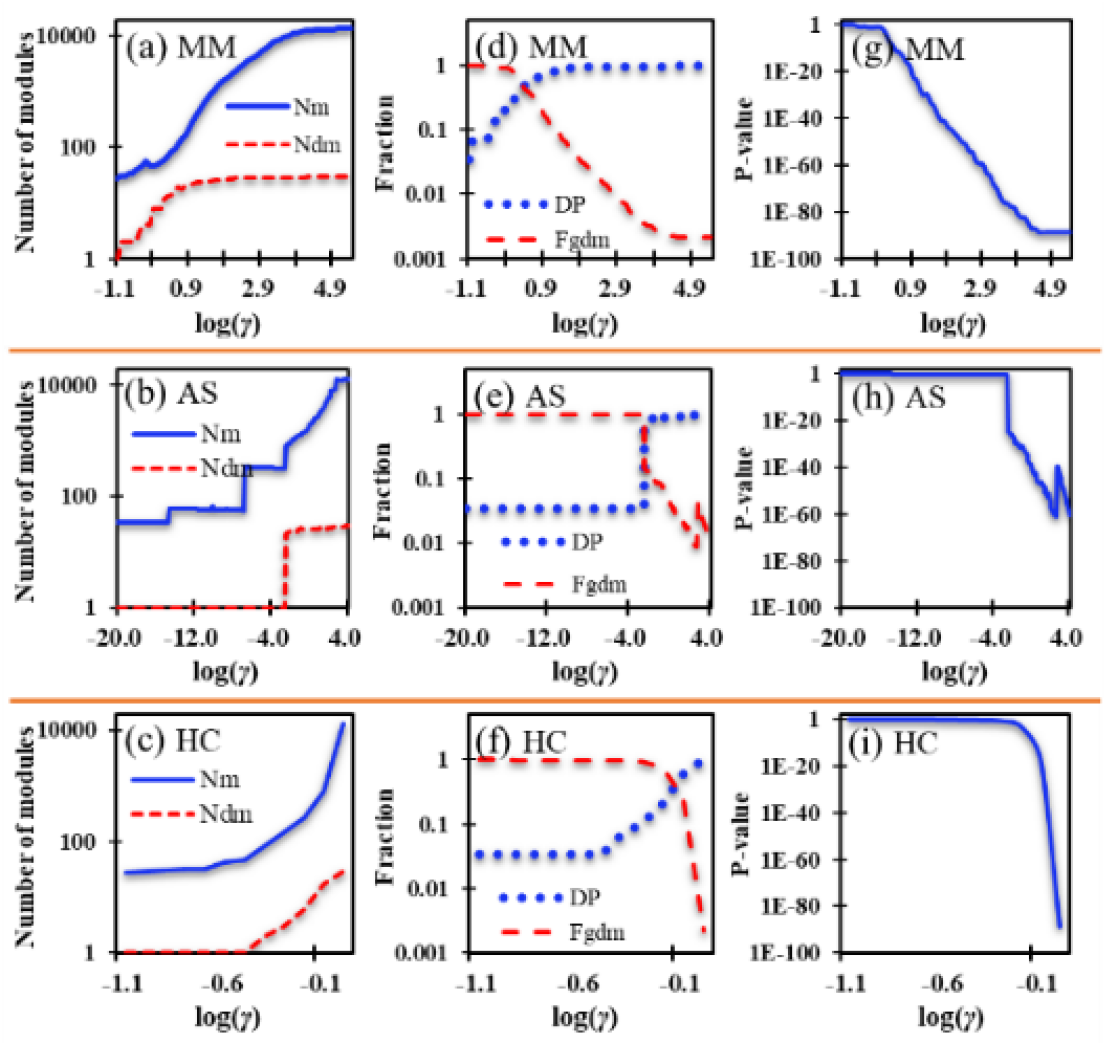
Statistics of modules and genes related to Alzheimer Disease. N_m_ and N_dm_ denotes the number of modules and the number of disease-related modules (i.e., modules containing disease genes), respectively. DP indicates the degree of dispersion of disease-related genes, which is defined as the ratio of the number of disease-related modules to its maximum (i.e., the number of known disease genes). F_gdm_ denotes the fraction of genes belonging to disease-related modules among all genes. P-value measures the enrichment of disease genes in disease-related modules. (a)- (c) the N_m_- and N_dm_-values as a function of *γ*-values for three multiscale algorithms (MM, AS and HC). (d)-(f) the DP and F_gdm_ as a function of *γ*-values for the three algorithms. (g)-(i) the P-value as a function of *γ*-values for the three algorithms.

### Supplementary Note 3: Cross-validation in PPI network and disease-gene heterogeneous network

The eight baseline algorithms are used to study the performance of HyMM:

1. Four baseline algorithms in the PPI network: RWR (Köhler, et al., 2008), KS (Chen, et al., 2009), VS(Zhu, et al., 2012) and ICN (Hsu, et al., 2011);
2. Four baseline algorithms in the disease-gene heterogeneous network: RWRH (Li and Patra, 2010), CIPHER (Wu, et al., 2008), BiRW (Xie, et al., 2015) and KATZ (Singh-Blom, et al., 2013).

**Figure S 3** and **Figure S 4** supplement the results of top-k recall curves of HyMM using AS and HC in the PPI network.

**Figure S 5** and **Figure S 6** supplement the results of top-k recall curves of HyMM using AS and HC in the disease-gene heterogeneous network.

**Figure S 3.**
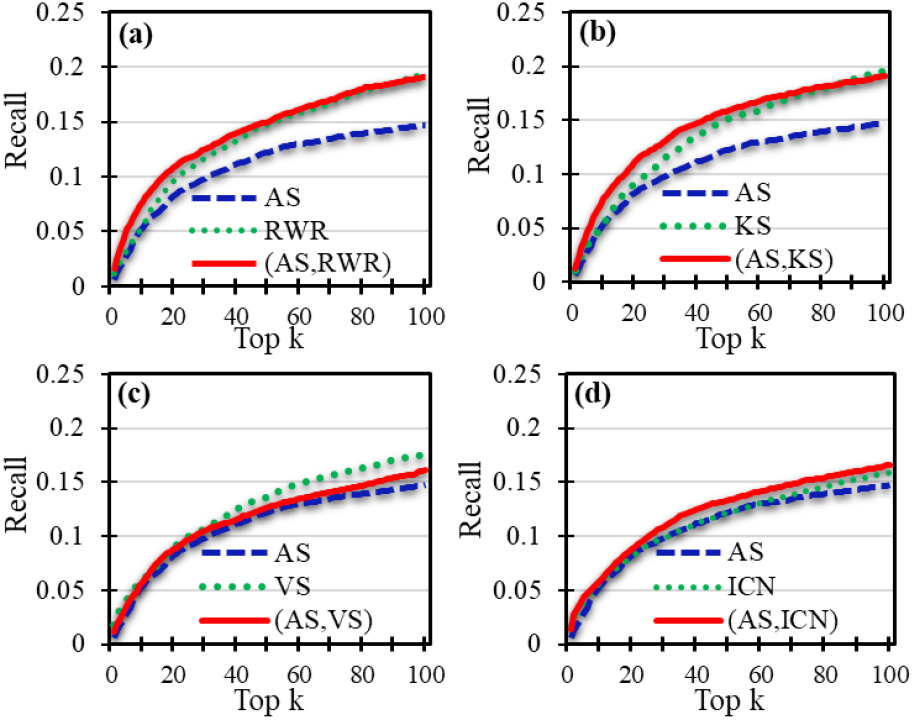
In the PPI network, top-k recall curves of HyMM algorithms **using AS**, compared to corresponding algorithms: (a) RWR, (b) KS, (c) VS, (d) ICN.

**Figure S 4.**
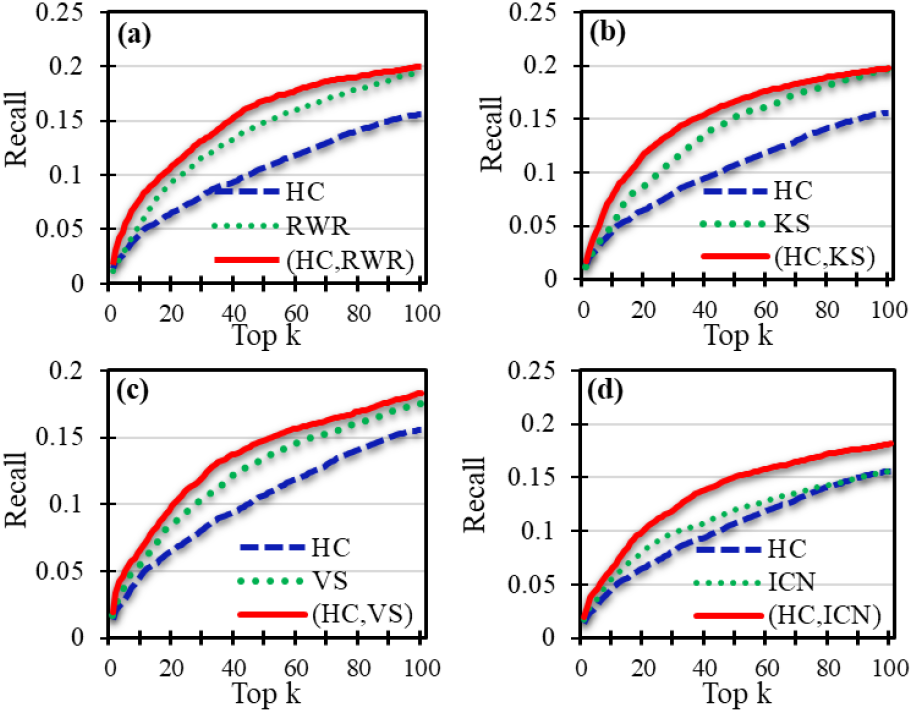
In the PPI network, top-k recall curves of HyMM algorithms **using HC**, compared to corresponding algorithms: (a) RWR, (b) KS, (c) VS, (d) ICN.

**Figure S 5.**
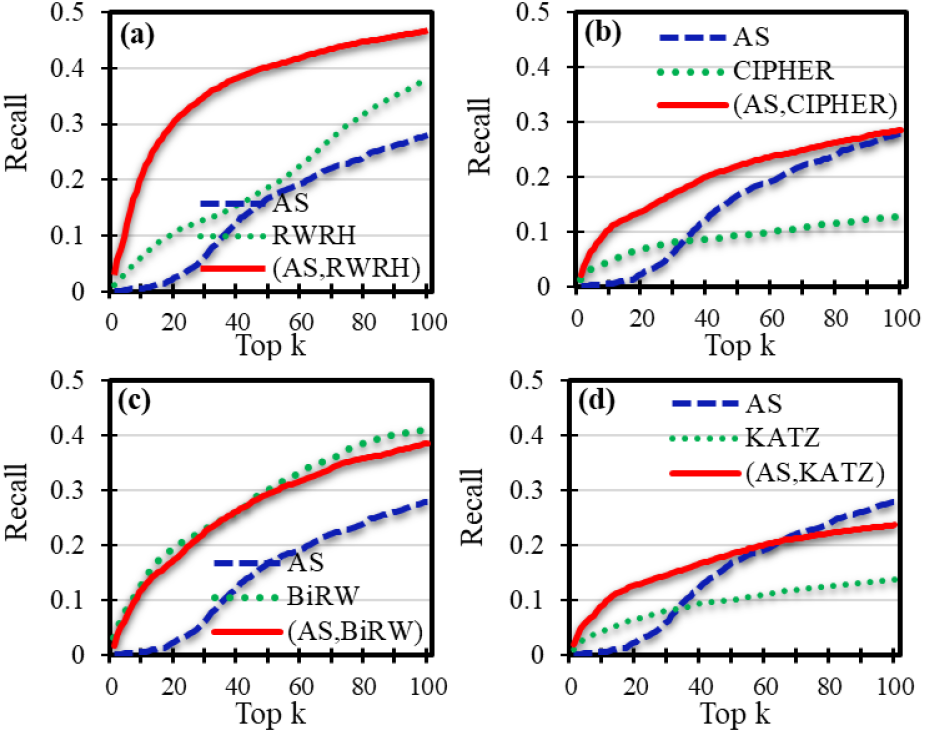
In the disease-gene heterogeneous network, top-k recall curves of HyMM algorithms **using AS**, compared to corresponding baseline algorithms: (a) RWRH, (b) CIPHER, (c) BiRW, (d) KATZ.

**Figure S 6.**
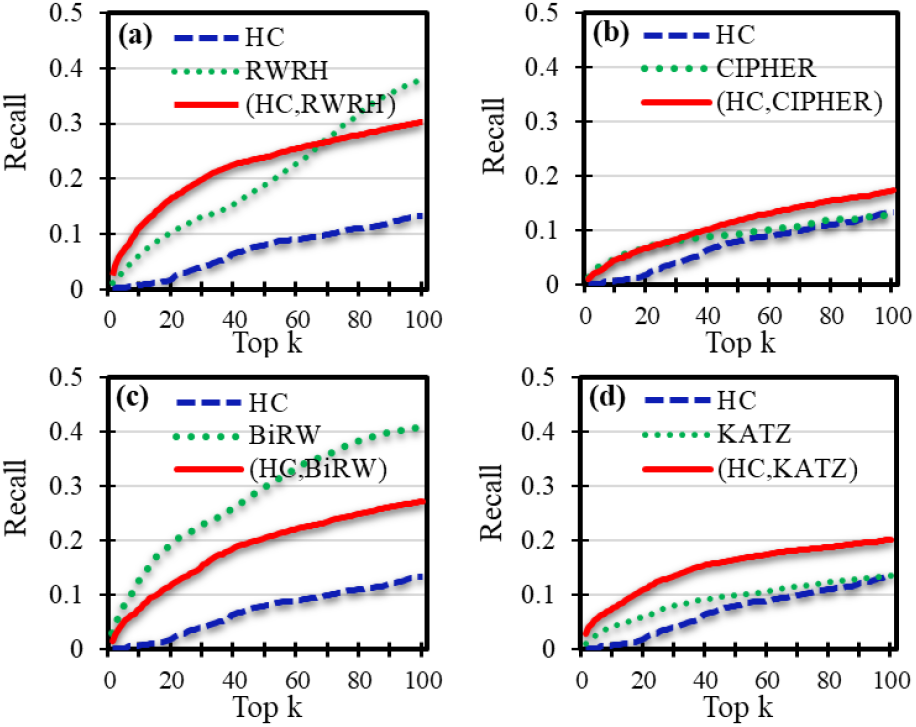
In the disease-gene heterogeneous network, top-k recall curves of HyMM algorithms **using HC**, compared to corresponding baseline algorithms: (a) RWRH, (b) CIPHER, (c) BiRW, (d) KATZ.

### Supplementary Note 4: Performance evaluation on external dataset

Here, we display the evaluation results of HyMM and baseline algorithms on external dataset (i.e., DisGeNet), also called IndTest. In this case, we calculate the scores of candidate genes by using the Mesh disease-gene associations as training set, and then evaluate the prediction performance by using the disease-gene associations belonging only to DisGeNet as test set.

**Figure S 7.**
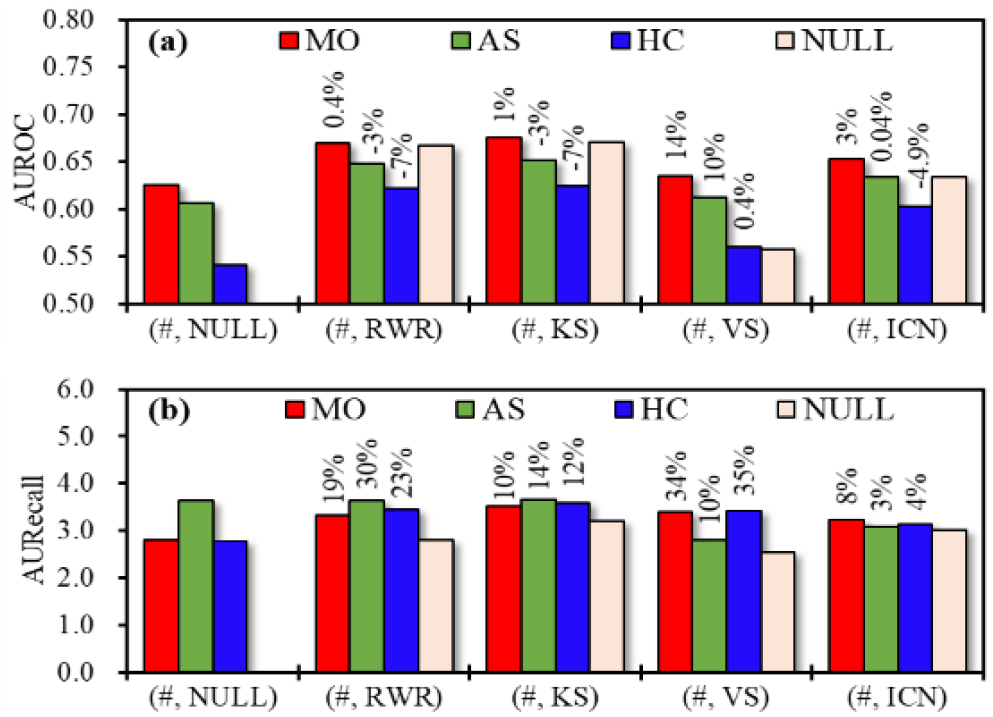
In the PPI network, for IndTest, performance comparison of (a) AUROC and (b) AURecall obtained by HyMM algorithms and four baseline algorithms. Multiscale module identification algorithms: MO, AS and HC. “(#, RWR)” denotes the HyMM algorithm integrating MO/AS/HC with RWR. “NULL” means that the multi-scale algorithm or baseline algorithm is not applicable. The percentage on the bar is performance improvement ratio to baseline algorithm.

**Figure S 8.**
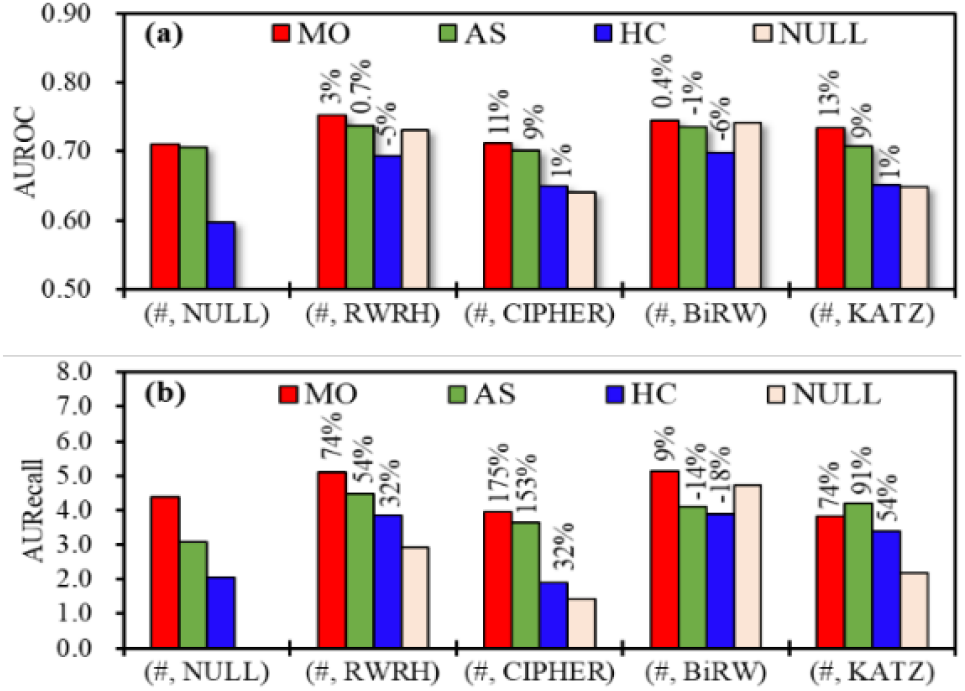
In the disease-gene heterogeneous network, for IndTest, performance comparison of (a) AUROC and (b) AURecall obtained by HyMM algorithms and four baseline algorithms. Multiscale module identification algorithms: MO, AS and HC. “(#, RWRH)” denotes the HyMM algorithm integrating MO/AS/HC with RWRH. “NULL” means that the multi-scale algorithm or baseline algorithm is not applicable. The percentage on the bar is performance improvement ratio to baseline algorithm.

### Supplementary Note 5: Case study

As an example, we obtain the top-20 genes from the ranking list of genes related to Alzheimer’s disease (AD). **Table S 1** shows the top-20 predicted genes for AD, the druggability of the candidate genes (Finan, et al., 2017; Freshour, et al., 2020; Wang, et al., 2019) as well as related information. **Table S 2** and **Table S 3** show the results of KEGG and GO enrichment analysis for the candidate genes.

Through literature verification, we find that this list of genes contains many genes experimentally proven to have associations with AD. For example, Park, et al. (2021) showed that ALK was important to the tau-mediated AD pathology; Annunziata, et al. (2013) showed that the deficiency of NEU1 caused the occurrence of an AD-like amyloidogenic process; Lian, et al. (2015) showed that the dysregulation of neuron-glia interaction through NFκB/C3/C3aR signaling may lead to synaptic dysfunction in AD; Rasmussen, et al. (2018) confirmed that the low baseline levels of complement C3 were associated with high risk of AD; Pichiah, et al. (2020) showed that C4B was differentially expressed in AD; Michele, et al. (2017) observed a statistically significant increase of CNVs for C4B in AD patients, suggesting a possible role for C4A CNVs in the risk of AD; Stoye, et al. (2020) demonstrated that APOA1 might be a key factor within intestine altered in AD-like pathology.

**Table S 1.**
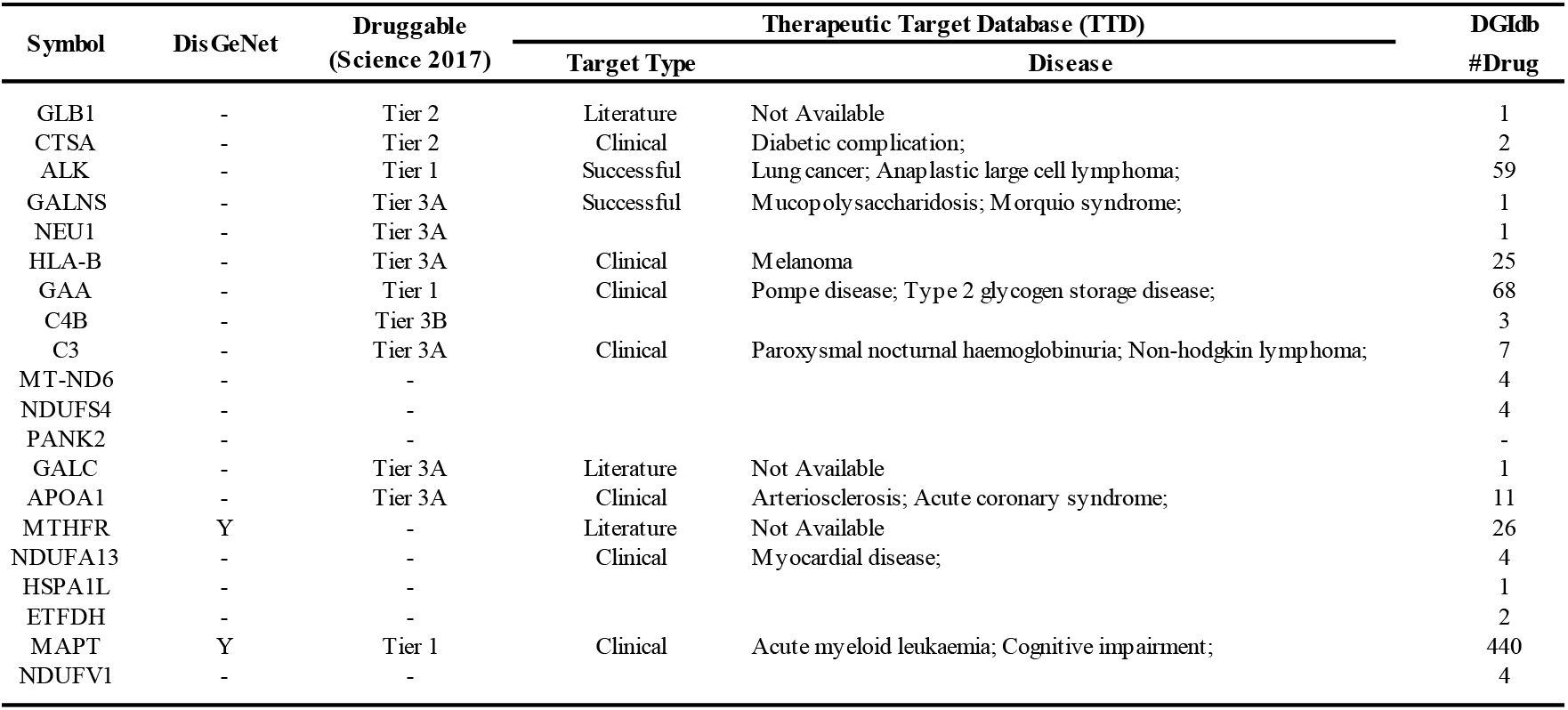
Analysis of candidate genes for AD. “Y” denotes that there exists record in DisGeNet. Tier 1 denotes genes that encode protein targets of approved or clinical trial-phase drug candidates and tier 2 denotes genes that encode protein targets with high sequence similarity to tier 1 proteins or targeted by small drug-like molecules.

**Table S 2.** Enrichment analysis of KEGG pathways for top-k predicted genes related to AD.

| #term ID | term description | FDR |
| --- | --- | --- |
| hsa04142 | Lysosome | 1.6E-07 |
| hsa01100 | Metabolic pathways | 3.6E-06 |
| hsa00190 | Oxidative phosphorylation | 1.9E-04 |
| hsa00600 | Sphingolipid metabolism | 2.0E-04 |
| hsa04723 | Retrograde endocannabinoid signaling | 2.0E-04 |
| hsa05012 | Parkinson's disease | 2.0E-04 |
| hsa05010 | Alzheimer's disease | 2.1E-04 |
| hsa04714 | Thermogenesis | 6.0E-04 |
| hsa00511 | Other glycan degradation | 1.2E-03 |
| hsa00531 | Glycosaminoglycan degradation | 1.2E-03 |
| hsa00052 | Galactose metabolism | 2.6E-03 |
| hsa04932 | Non-alcoholic fatty liver disease (NAFLD) | 2.6E-03 |
| hsa05016 | Huntington's disease | 4.6E-03 |
| hsa05150 | Staphylococcus aureus infection | 5.7E-03 |
| hsa05134 | Legionellosis | 5.9E-03 |
| hsa04612 | Antigen processing and presentation | 8.1E-03 |
| hsa05133 | Pertussis | 9.5E-03 |
| hsa04610 | Complement and coagulation cascades | 9.9E-03 |
| hsa05322 | Systemic lupus erythematosus | 1.4E-02 |
| hsa04145 | Phagosome | 2.9E-02 |
| hsa05167 | Kaposi's sarcoma-associated herpesvirus infection | 4.2E-02 |
| hsa05168 | Herpes simplex infection | 4.2E-02 |
| hsa05169 | Epstein-Barr virus infection | 4.2E-02 |
| hsa05203 | Viral carcinogenesis | 4.2E-02 |

**Table S 3.**
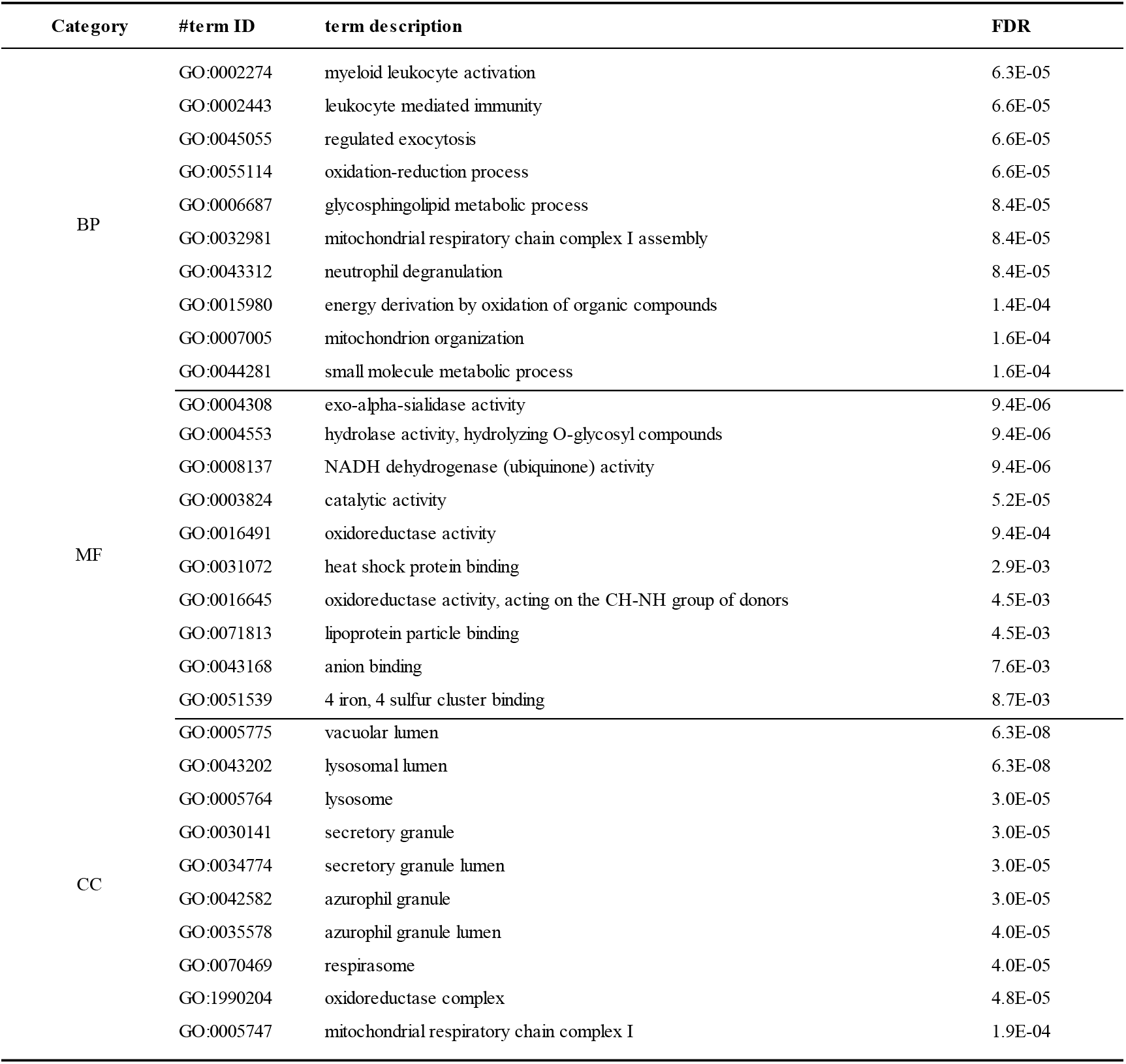
Enrichment analysis of GO for top-k predicted genes related to AD.

| Category | #term ID | term description | FDR |
| --- | --- | --- | --- |
| BP | GO:0002274 | myeloid leukocyte activation | 6.3E-05 |
|  | GO:0002443 | leukocyte mediated immunity | 6.6E-05 |
|  | GO:0045055 | regulated exocytosis | 6.6E-05 |
|  | GO:0055114 | oxidation-reduction process | 6.6E-05 |
|  | GO:0006687 | glycosphingolipid metabolic process | 8.4E-05 |
|  | GO:0032981 | mitochondrial respiratory chain complex I assembly | 8.4E-05 |
|  | GO:0043312 | neutrophil degranulation | 8.4E-05 |
|  | GO:0015980 | energy derivation by oxidation of organic compounds | 1.4E-04 |
|  | GO:0007005 | mitochondrion organization | 1.6E-04 |
|  | GO:0044281 | small molecule metabolic process | 1.6E-04 |
| MF | GO:0004308 | exo-alpha-sialidase activity | 9.4E-06 |
|  | GO:0004553 | hydrolase activity, hydrolyzing O-glycosyl compounds | 9.4E-06 |
|  | GO:0008137 | NADH dehydrogenase (ubiquinone) activity | 9.4E-06 |
|  | GO:0003824 | catalytic activity | 5.2E-05 |
|  | GO:0016491 | oxidoreductase activity | 9.4E-04 |
|  | GO:0031072 | heat shock protein binding | 2.9E-03 |
|  | GO:0016645 | oxidoreductase activity, acting on the CH-NH group of donors | 4.5E-03 |
|  | GO:0071813 | lipoprotein particle binding | 4.5E-03 |
|  | GO:0043168 | anion binding | 7.6E-03 |
|  | GO:0051539 | 4 iron, 4 sulfur cluster binding | 8.7E-03 |
| CC | GO:0005775 | vacuolar lumen | 6.3E-08 |
|  | GO:0043202 | lysosomal lumen | 6.3E-08 |
|  | GO:0005764 | lysosome | 3.0E-05 |
|  | GO:0030141 | secretory granule | 3.0E-05 |
|  | GO:0034774 | secretory granule lumen | 3.0E-05 |
|  | GO:0042582 | azurophil granule | 3.0E-05 |
|  | GO:0035578 | azurophil granule lumen | 4.0E-05 |
|  | GO:0070469 | respirasome | 4.0E-05 |
|  | GO:1990204 | oxidoreductase complex | 4.8E-05 |
|  | GO:0005747 | mitochondrial respiratory chain complex I | 1.9E-04 |

### Supplementary Note 6: Parameter stability

We here display the effect of the sampling parameter *Δ log γ* on prediction performance. We have calculated the AUROC and AURecall values of all HyMM algorithms in the interval of *Δ log γ* ∈ [0.01,0.2]. The results show that the HyMM algorithms using MO and AS are very stable to the sampling parameter, while the HyMM algorithms using HC have relatively large fluctuations.

**Figure S 9.**
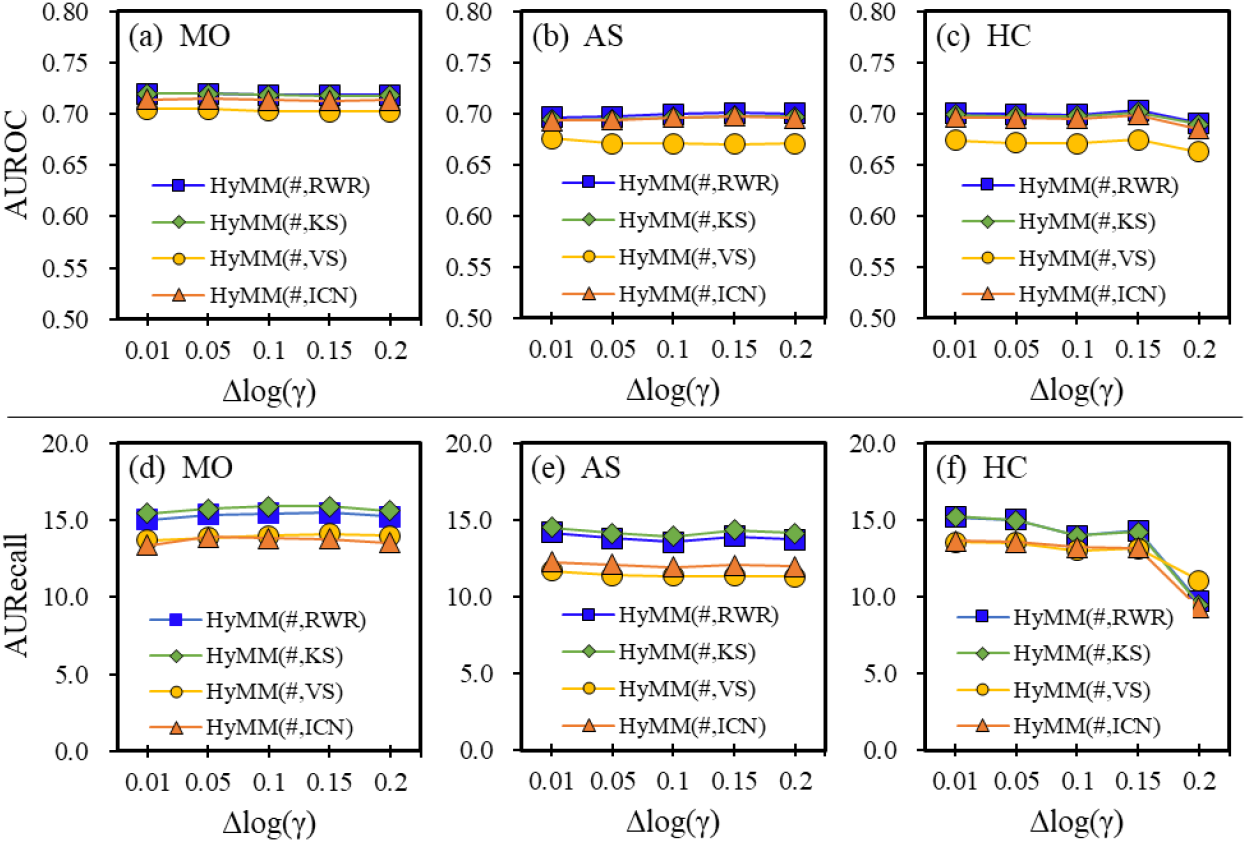
In the PPI network, (a)-(c) Effect of resolution interval on the AUROC performance of HyMM using MO, AS and HC respectively; (d)-(f) Effect of resolution interval on the AURecall performance of HyMM using MO, AS and HC respectively.

**Figure S 10.**
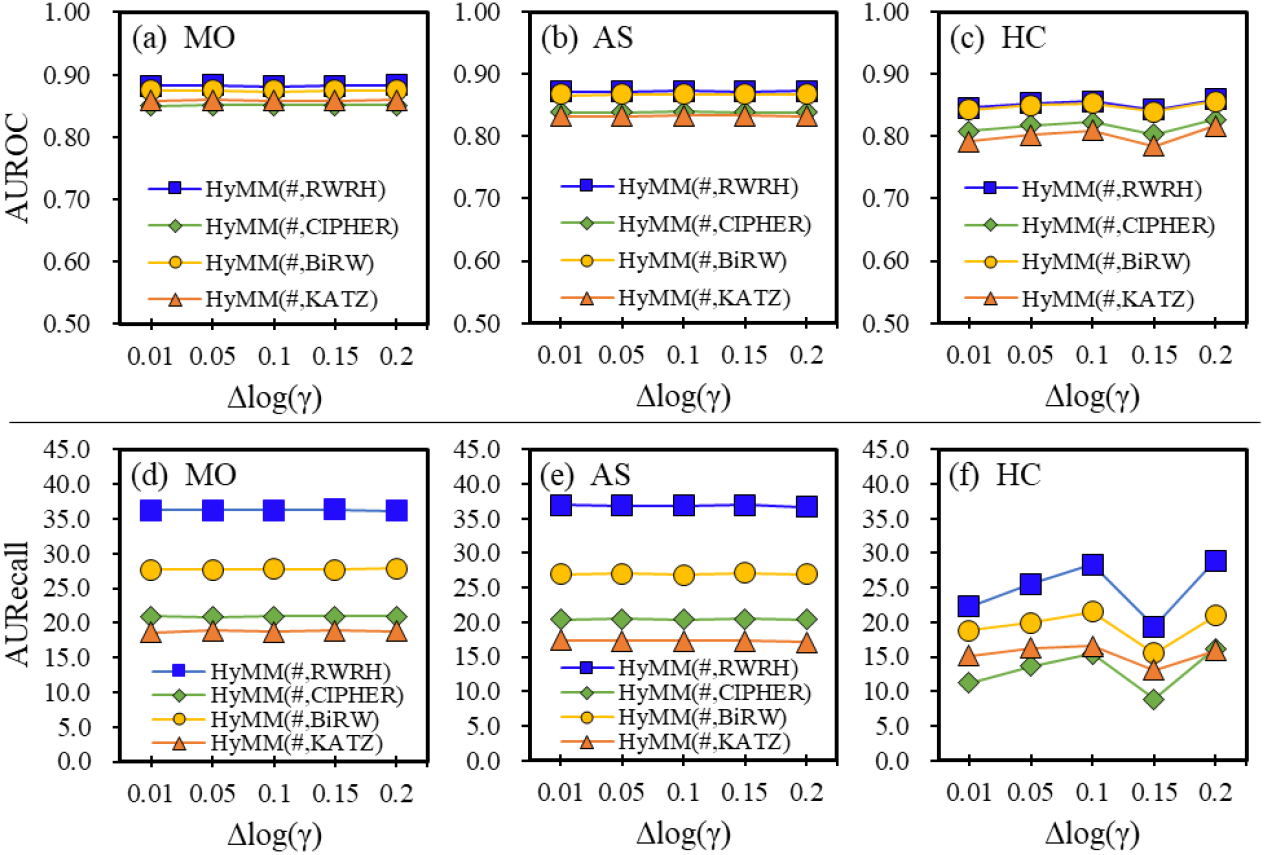
In the disease-gene heterogeneous network, (a)-(c) Effect of resolution interval on the AUROC performance of HyMM using MO, AS and HC respectively; (d)-(f) Effect of resolution interval on the AURecall performance of HyMM using MO, AS and HC respectively.

## Notes

https://github.com/xiangju0208/HyMM

### Competing Interest Statement

The authors have declared no competing interest.

